# CoMR: an integrative scoring pipeline for Comprehensive Mitochondrial proteome Reconstruction across eukaryotes

**DOI:** 10.64898/2026.02.20.707009

**Authors:** Julie Boisard, Shelby K. Williams, Andrew J. Roger, Courtney W. Stairs

## Abstract

Mitochondrial proteome reconstruction from eukaryotic sequence data typically relies on prediction of mitochondrial targeting signals (MTSs). However, MTS predictors are primarily trained on model organisms and may perform poorly in phylogenetically divergent lineages or in organisms with atypical or reduced targeting sequences. Accurate reconstruction therefore requires integration of complementary sources of evidence beyond targeting prediction alone. We developed CoMR (Comprehensive Mitochondrial Reconstructor), an integrative workflow that combines targeting prediction, curated homology searches, large-scale similarity searches, and automated phylogenetic analysis within a unified scoring framework. Benchmarking on the model yeast *Saccharomyces cerevisiae* yielded strong discriminatory performance (ROC-AUC = 0.92), exceeding standalone TargetP2 prediction (ROC-AUC = 0.72). In the divergent anaerobic protist *Paratrimastix pyriformis*, CoMR maintained robust performance (ROC-AUC = 0.86) validated with an experimental proteome despite extreme class imbalance, achieving a precision-recall AUC of 0.183 (∼78-fold enrichment over random expectation and ∼10-fold improvement over TargetP2). Ablation analyses demonstrate that predictive performance is robust to individual evidence-layer removal and that the relative contribution of homology sources depends on phylogenetic context. Together, these results show that integrative evidence scoring improves mitochondrial proteome reconstruction across both model and non-model eukaryotes.

## Introduction

Mitochondria are central to eukaryotic cellular metabolism, participating in ATP production through oxidative phosphorylation as well as amino acid metabolism, iron-sulfur cluster biogenesis, and organellar genome maintenance^1–3^. In well-studied aerobic organisms such as humans and yeast, mitochondrial proteomes have been extensively characterized, providing detailed insight into organellar composition and function^4^. However, accurate reconstruction of mitochondrial proteomes remains challenging in many eukaryotic lineages, particularly those that are phylogenetically divergent or adapted to anaerobic environments.

Anaerobic protists represent some of the most understudied yet phylogenetically diverse branches of the eukaryotic tree of life. Many possess mitochondrion-related organelles (MROs), which are highly reduced versions of canonical mitochondria^5,6^. MROs vary in structure and metabolic capacity across lineages, often exhibiting lineage-specific adaptations and reductions^7^. Studying these organelles provides insight into mitochondrial evolution and organelle plasticity, but reconstructing their proteomes is computationally and experimentally challenging.

In silico prediction of mitochondrial localization typically relies on detection of N-terminal mitochondrial targeting signals (MTSs). Most widely used MTS predictors are trained on model organisms and assume canonical targeting sequence properties. In divergent lineages such as metamonads, MTSs can be highly atypical or absent, and protein import may rely more heavily on internal targeting elements^8–10^. Consequently, targeting-based approaches alone may underestimate mitochondrial proteome size or misclassify divergent proteins. Homology-based searches can complement targeting prediction, but database composition and phylogenetic representation strongly influence performance. A unified framework that integrates multiple independent sources of evidence for mitochondrial localization is therefore needed.

Here, we present CoMR (Comprehensive Mitochondrial Reconstructor), an integrative workflow for mitochondrial and MRO proteome reconstruction. CoMR combines targeting prediction, curated and large-scale homology searches, profile-based HMM detection, and automated phylogenetic analysis within a unified scoring framework. By integrating partially independent evidence layers, CoMR generates a ranked prediction of mitochondrial localization that is adaptable to diverse phylogenetic contexts. We benchmark CoMR on experimental dataset from both a model organism (*Saccharomyces cerevisiae*) and a divergent anaerobic protist (*Paratrimastix pyriformis*), demonstrating improved predictive performance over standalone targeting prediction and robust behavior under extreme class imbalance.

## Methods

### Overview of the CoMR workflow

CoMR (Comprehensive Mitochondrial Reconstructor) is a modular Snakemake-based pipeline designed to identify and score mitochondrial and mitochondrial-related organelle (MRO) proteins from predicted eukaryotic proteomes. The workflow integrates targeting signal predictors, profile hidden Markov models (HMMs), homology searches, and phylogenetic validation into a unified evidence framework that produces ranked candidate lists suitable for expert curation (Figure 1).

**Figure 1.**
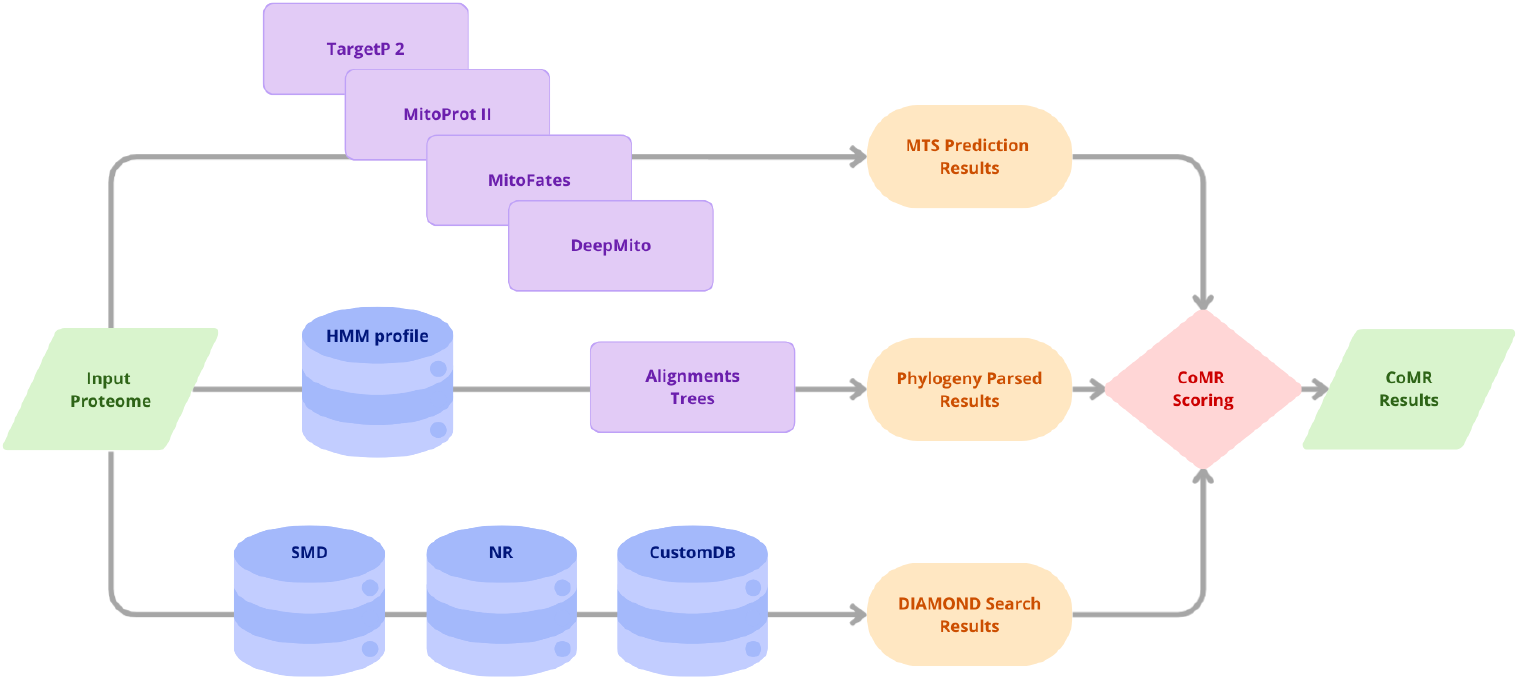
Overview of the CoMR pipeline. CoMR integrates targeting signal prediction, homology searches, and phylogenetic analysis to reconstruct a putative mitochondrial or mitochondrial-related organelle (MRO) proteome from a predicted proteome (Input Proteome). Three complementary evidence streams are computed in parallel. MTS prediction branch: TargetP2, MitoProt, MitoFates, and DeepMito predict mitochondrial targeting signals; their outputs are aggregated as MTS targeting prediction results. Phylogenetic branch: HMM profile searches identify candidate protein families, followed by phylogenetic reconstruction and automated tree parsing to classify sequences as mitochondrial or non-mitochondrial. Sequence similarity branch: Subtractive Mitochondrial Database (SMD), NR, and CustomDB. DIAMOND searches provide homology-based evidence, summarized as DIAMOND search results. All intermediate evidence layers are integrated by a user-defined scoring scheme to produce the final CoMR scores, which summarize mitochondrial support across all evidence categories.

### Input data and preprocessing

CoMR accepts one or more predicted proteomes in FASTA format. The input protein sequences can be derived from gene models, transcriptomes, or proteomic analysis. Each input FASTA undergoes normalization, during which sequence headers are replaced by unique identifiers to ensure compatibility with downstream tools and to enable reproducible joins across evidence sources. A mapping table linking original headers to internal identifiers is retained. A filtered FASTA containing only methionine-initiated sequences is generated for N-terminal targeting prediction steps, while the full indexed FASTA is preserved for homology-based analyses.

### Targeting and localization prediction

Four independent predictors are applied to each normalized proteome to infer mitochondrial targeting signals and subcellular localization. TargetP 2.0^11^ is used to predict mitochondrial targeting peptides and cleavage sites, while MitoFates^12^ and Mitoprot II^13^ provide complementary estimates of presequence presence, export probability, and cleavage positions. DeepMito^14^, a deep-learning-based classifier, is run using a bundled UniProt^15^ reference FASTA to infer localization probabilities. Raw outputs from each predictor are retained, and parsed summary tables are generated to standardize scores and categorical calls across tools.

### Homology and profile-based searches

DIAMOND^16^ BLASTP searches are executed against multiple protein databases, including the CoMR Subtractive Mitochondrial Database (SMD). The SMD is derived from the proteome of six organisms (*Andalucia godoyi*^17^, *Tetrahymena thermophila*^18,19^, *Arabidopsis thaliana*^20^, *Homo sapiens*^21^, *Acanthamoeba castellanii*^22,23^, *Saccharomyces cerevisiae*^24^) divided into two fasta files with one file containing the sequences of curated mitochondrial proteins and the other file containing sequences of non-mitochondrial proteins. To capture conserved mitochondrial protein families, CoMR performs profile HMM searches using HMMER^25^ (hmmscan) using a curated CoMR mitochondrial profile HMM database derived from the SMD using default settings. Significant hits are parsed and used to nominate sequences for downstream alignment and phylogenetic analysis. Input sequences can optionally be used as queries agains the NCBI non-redundant (NR) database: when NR searches are enabled, taxonomy annotation is reconstructed using NCBI taxonomy dump files^26^. An optional CustomDB search can be activated to use the input proteomes as queries against a user-provided DIAMOND database for lineage-specific analyses. The construction and usage of the SMD is detailed in the Supplementary Data (Supplementary Methods - CoMR databases), along with a description of its usage in CoMR’s profile HMM database, phylogenetic analysis and tree parsing function. All CoMR databases are available on the CoMR Figshare repository (https://doi.org/10.17044/scilifelab.31361839).

### Alignment and phylogenetic inference

Sequences with significant HMM hits are incorporated into reference multiple sequence alignments using MAFFT 7^27^, followed by automated trimming with trimAl to remove poorly aligned regions. For each trimmed alignment, phylogenetic trees are inferred using IQ-TREE 2^28^ under an LG+G model with fast heuristics enabled. Trees are parsed with ete3^29^, by rooting each tree on the query sequence and propagating mitochondrial/other states via a Fitch^30^-style traversal: each query sequence is classified based on its phylogenetic relationship to known mitochondrial and non-mitochondrial reference sequences.

### Evidence integration and scoring

All parsed evidence layers, including targeting predictions, HMMscan hits, blast homology scores and taxonomy, and phylogenetic classifications, are merged into a single per-sequence table. CoMR applies a configurable scoring rubric defined in external scorecard files, allowing flexible weighting of individual evidence sources.

CoMR integrates heterogeneous evidence sources at the level of individual protein sequences using a rule-based scoring framework. Each layer of evidence provides a clue as to the presence or absence of support for mitochondrial localization. Under the default scoring scheme, each qualifying evidence source contributes equally to a cumulative CoMR score, which is reported alongside detailed per-component annotations to ensure interpretability and traceability. A sequence receives one point for each mitochondrial targeting predictor reporting a positive prediction (TargetP, MitoProt, or MitoFates), one point for curated homology support based on matches to multiple mitochondrial datasets in the SMD, one point for phylogenetic placement within a mitochondrial clade, and one point for large-scale homology support from the NCBI non-redundant database when hits are annotated with mitochondrion-related keywords. DeepMito predictions are not included in the composite score, as this tool assumes mitochondrial localization and instead provides sub-mitochondrial localization information. DeepMito outputs are reported alongside the composite score to facilitate downstream interpretation. The default equal-scoring scheme assigns equal weight to all contributing evidence layers, resulting in a composite score ranging from 0 to 6. Alternative scoring schemes can be defined by modifying the scorecard file provided with CoMR. The detailed binarization rules and scoring logic are described in Supplementary Data (Supplementary Methods - CoMR evidence integration and scoring).

### Workflow execution and containerization

CoMR is distributed as a fully containerized workflow to ensure reproducibility across computing environments. CoMR’s Docker^31^ image and Singularity/Apptainer^32^ image can be retrieved from the CoMR GitHub repository (https://github.com/theLabUpstairs/CoMR). Licensed software (TargetP2) and large external databases are mounted at runtime rather than bundled in the image. The workflow is executed via Snakemake within the container, with user-configurable resource limits for parallelization and memory-intensive steps such as DIAMOND searches.

### Software environment

The runtime environment is defined via a Conda specification in the container and includes Python 3.11, Snakemake^33^, HMMER, DIAMOND, MAFFT, IQ-TREE 2, Biopython^34^, pandas, and supporting libraries required for parsing, scoring, and phylogenetic analysis.

### CoMR performance evaluation

Benchmarking was performed on one model organism (*Saccharomyces cerevisiae*) and one non-model organism (*Paratrimastix pyriformis*). To prevent circularity, benchmarking-specific versions of all internal reference databases were generated by excluding sequences from the evaluated species prior to homology searches. In addition, DIAMOND searches against the NCBI non-redundant (NR) database were parsed using a taxonomically controlled exclusion strategy, in which hits assigned to the evaluated species or higher taxonomic ranks were removed prior to evidence integration. Performance was assessed using threshold-based metrics, ROC-AUC and precision-recall analyses across CoMR score thresholds (0-6) and compared to individual mitochondrial targeting predictors using their recommended cutoffs. Detailed benchmarking is described in Supplementary Data (Supplementary Methods - CoMR performance evaluation) and available on the CoMR Figshare repository (https://doi.org/10.17044/scilifelab.31361839).

## Results

### Overview of CoMR outputs and composite scoring

For each input proteome, CoMR produces a unified evidence table in which all targeting predictions, homology-based signals, and phylogenetic classifications are integrated at the level of individual protein sequences. Each sequence is assigned a composite CoMR score summarizing the number of independent lines of evidence supporting mitochondrial localization. The output table reports, for each sequence, the composite CoMR score together with the individual contributions of targeting predictors, homology searches, phylogenetic classification, and associated annotation fields. Under the default “equal” scoring scheme, the CoMR score ranges from 0 to 6, with higher values indicating increasing support from multiple evidence layers, including mitochondrial targeting predictions, curated homology matches, large-scale homology searches, and phylogenetic placement. DeepMito predictions are reported alongside the score to provide sub-mitochondrial localization information but are not included in the composite score. The resulting score provides a continuous ranking of candidate proteins rather than a binary classification, allowing users to apply dataset-specific thresholds or to prioritize high-scoring sequences for downstream analyses and mitochondrial proteome reconstruction. A schematic overview of the CoMR workflow is shown in Figure 1.

### Benchmarking strategy and evaluation framework

To evaluate the performance of CoMR, we benchmarked its ability to predict mitochondrial localization using both a well-annotated model organism (*Saccharomyces cerevisiae*) and a non-model metamonad (*Paratrimastix pyriformis*). This design allowed us to assess CoMR performance under conditions of extensive reference data availability as well as in a lineage with highly reduced and divergent mitochondrial-related organelles. To prevent circularity during benchmarking on *S. cerevisiae*, all internal reference databases used by CoMR were reconstructed after excluding *S. cerevisiae* sequences, as described in the Supplementary Data (Supplementary Methods - CoMR performance evaluation). In addition, homology evidence derived from the NCBI non-redundant (NR) database was controlled by taxonomic exclusion of hits assigned to *S. cerevisiae* or its close relatives. Benchmarking was performed under multiple exclusion stringencies (species-, genus-, and order-level), with stricter conditions reported in the Supplementary Data (Supplementary Methods - CoMR performance evaluation, Supplementary Table S1, Supplementary Figure S1).

For *Paratrimastix pyriformis*, benchmarking was performed using both the whole-cell proteome and a curated mitochondrial-related organelle (MRO) proteome^35^, with taxonomic exclusion of its NCBI taxonomic identifier from NR-derived evidence. CoMR performance was evaluated using the default equal-weight scoring scheme, in which all six evidence layers contribute equally to the composite score (range 0-6). For each dataset, true positive rate (TPR) and false positive rate (FPR) were calculated across all score thresholds and used to construct receiver operating characteristic (ROC) curves and corresponding area under the curve (ROC-AUC) values. CoMR performance was compared to the baseline performance of TargetP2 using identical reference sets. Because mitochondrial proteins represent a small fraction of the *P. pyriformis* proteome, precision-recall (PR) curves and area under the PR curve (AUPR) values were additionally computed for this dataset.

### CoMR performance on a model organism, *Saccharomyces cerevisiae*

CoMR was applied to the *S. cerevisiae* proteome to evaluate its ability to distinguish mitochondrial from non-mitochondrial proteins. CoMR scores were compared to the curated reference proteome^24^, and true positive rate (TPR) and false positive rate (FPR) values were calculated across all possible score thresholds (0-6). These values were used to construct a ROC curve for the default equal-weight scoring scheme (Figure 2A). The resulting area under the ROC curve (ROC-AUC) was 0.92, indicating strong discriminatory performance of CoMR on the *S. cerevisiae* proteome.

**Figure 2.**
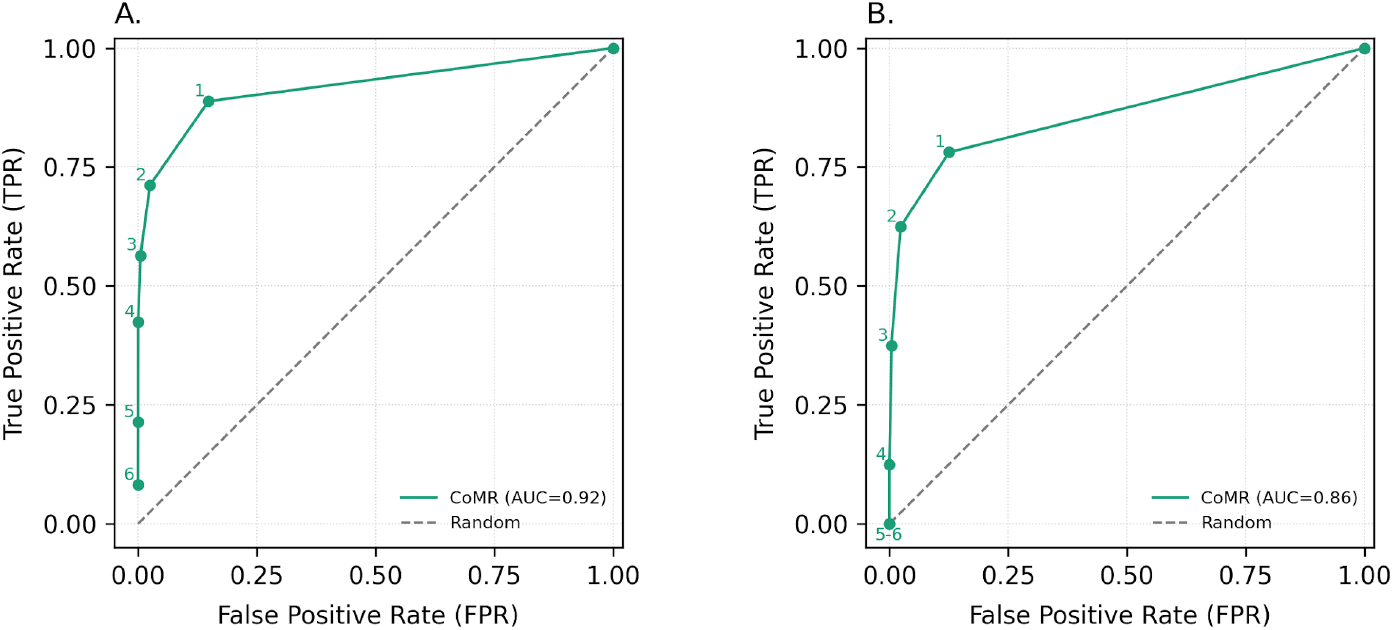
The threshold-resolved ROC curves for *S. cerevisiae* proteome (AUC = 0.92) and *P. pyriformis* proteome (AUC = 0.86) indicate strong performance, with CoMR maintaining high sensitivity at low false-positive rates across score cutoffs. **(A)** *S. cerevisiae* proteome. **(B)** *P. pyriformis* proteome. The ROC curves were computed using the equal scoring scheme, with possible scores ranging between 0 and 6 (noted next to data points). Note that scores 5 and 6 for *P. pyriformis* share the same data point. The dashed line represents the ROC curve for a classifier that randomly assigns positive and negative classes.

We first examined the baseline performance of the three mitochondrial targeting sequence (MTS) predictors on the *S. cerevisiae* proteome by comparing true positive (TP), false positive (FP), true negative (TN), and false negative (FN) counts for each predictor. All three predictors showed strong ability to correctly identify non-mitochondrial proteins, with relatively low FN rates (Figure 3A). However, their precision differed, with MitoProt producing a higher number of FP compared to TargetP2 and MitoFates. To assess the impact of this behavior on CoMR performance, we evaluated alternative scoring schemes in which MitoProt was either excluded entirely or downweighed by reducing its contribution to the overall score (Figure 4A). These modified schemes allowed us to determine whether MitoProt-associated FP disproportionately influenced CoMR’s predictive accuracy.

**Figure 3.**
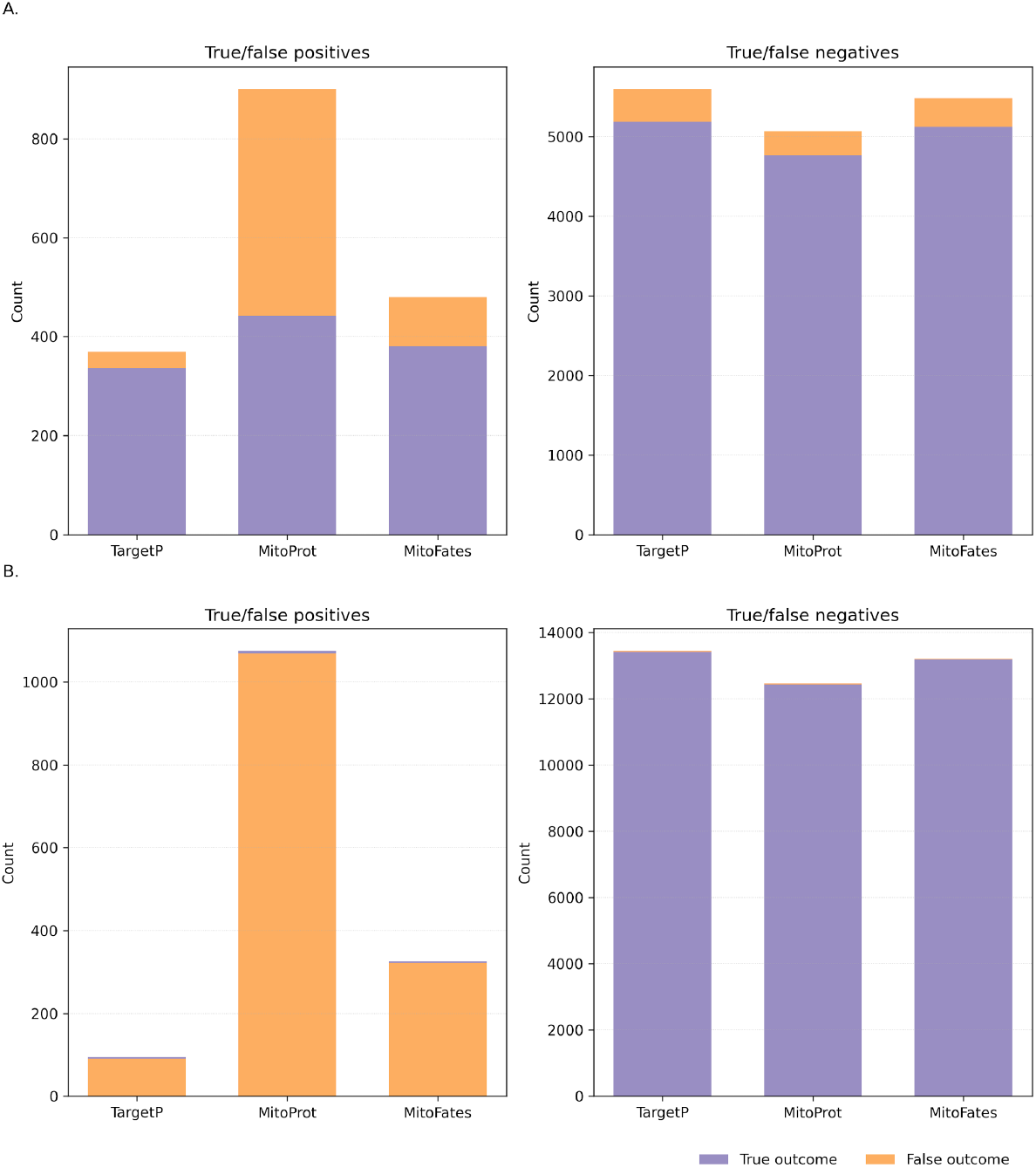
MTS predictor performance in the *S. cerevisiae* proteome and *P. pyriformis* proteome. **(A)** *S. cerevisiae* proteome. **(B)** *P. pyriformis* proteome. Stacked bar plots compare confusion counts for TargetP, MitoProt, and MitoFates at their CoMR recommended thresholds. Panel A shows true and false positives; panel B shows true and false negatives. True outcomes are shown in purple and false outcomes in orange.

**Figure 4.**
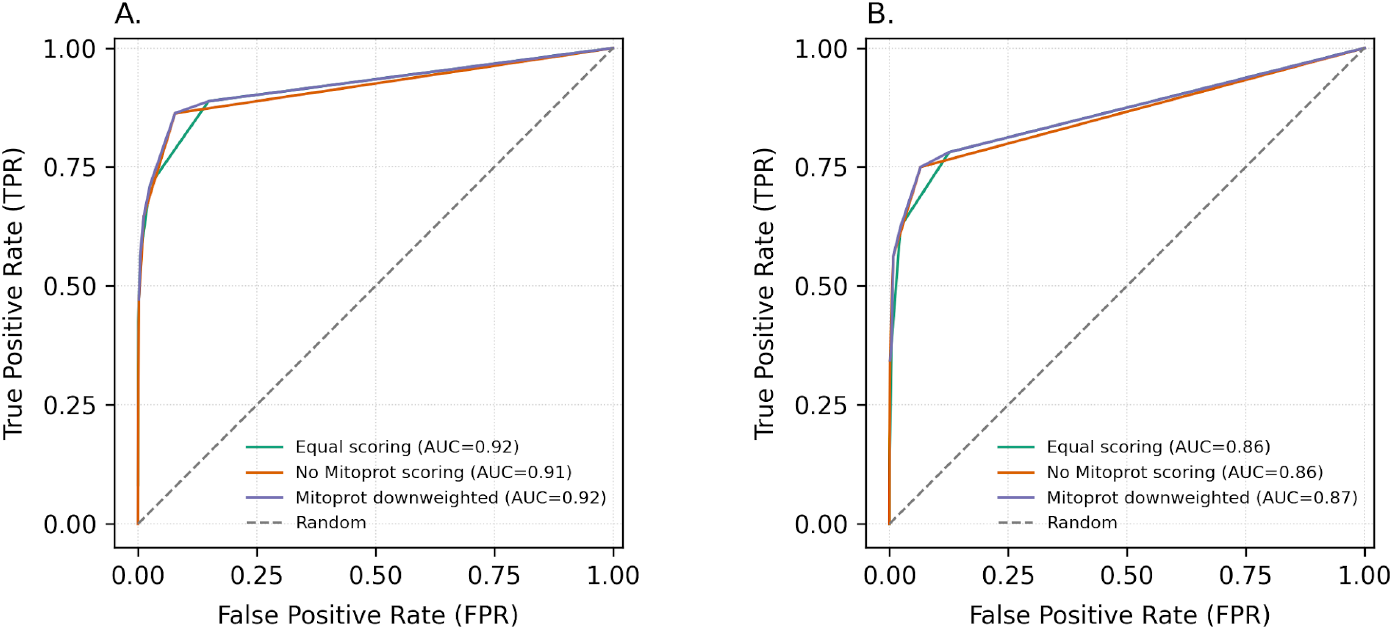
Mitoprot weighting slightly influences CoMR ROC performance in the *S. cerevisiae* proteome and *P. pyriformis* proteome. **(A)** *S. cerevisiae* proteome. **(B)** *P. pyriformis* proteome. ROC curves were computed under three Mitoprot-related scoring schemes: Equal scoring (turquoise), No Mitoprot scoring (orange), and Mitoprot downweighted (purple). Each curve is annotated with its AUC; the dashed line indicates a random classifier.

We then recalculated TPR and FPR values under these modified scoring schemes and reconstructed the corresponding ROC curves and ROC-AUC values (Figure 4A, Table 1). Down-weighting MitoProt (0.5 instead of 1 point) slightly improved ROC-AUC, whereas complete removal of MitoProt led to a modest decrease in performance relative to the default equal-weight scheme. These results indicate that although MitoProt contributes FP, its inclusion provides a complementary signal that supports overall predictive performance. The modest effect of weighting adjustments further suggests that CoMR performance is not dominated by any single evidence layer. Moreover, the scoring framework is implemented as a customizable scorecard, allowing users to adapt component weights and fine-tune CoMR to maximise performance according to the characteristics of their own datasets.

**Table 1.** AUC scores of ROC curves summarizing CoMR performance on *S. cerevisiae* and *P. pyriformis* proteomes, according to different scoring schemes.

| Scoring scheme ROC | *S. cerevisiae* AUC score | *P. pyrifformis* AUC score |
| --- | --- | --- |
| Equal scoring | 0.91782 | 0.86113 |
| No Mitoprot scoring | 0.91482 | 0.85875 |
| Mitoprot downweighted | 0.92202 | 0.86571 |
| No tree search scoring | 0.91074 | 0.84789 |
| No subtractive database scoring | 0.91285 | 0.83138 |
| No TargetP2 scoring | 0.91736 | 0.86220 |
| No Mitofates scoring | 0.91613 | 0.86215 |
| No "mito" nr search scoring | 0.85265 | 0.82215 |

To quantify the contribution of individual evidence layers, we systematically removed each component from the scoring scheme and recalculated ROC-AUC values for the *S. cerevisiae* proteome (Table 1; Figure 5A). This ablation analysis allowed us to assess the relative influence of each search strategy on overall predictive performance. Removal of NR-based homology support produced the largest decrease in ROC-AUC, indicating that large-scale homology searches contribute substantially to mitochondrial protein identification in *S. cerevisiae*. Performance remained stable even under stricter genus- and order-level NR exclusion (Supplementary Figure S1, Supplementary Table S1), indicating that CoMR performance was not solely dependent on near-identical homologs. In contrast, removal of the remaining evidence layers resulted in more moderate reductions in performance, suggesting that these components provide complementary and partially overlapping signals. This effect likely reflects the dense phylogenetic representation of fungal sequences in NR, which enhances the homology-based signal for *S. cerevisiae* proteins.

**Figure 5.**
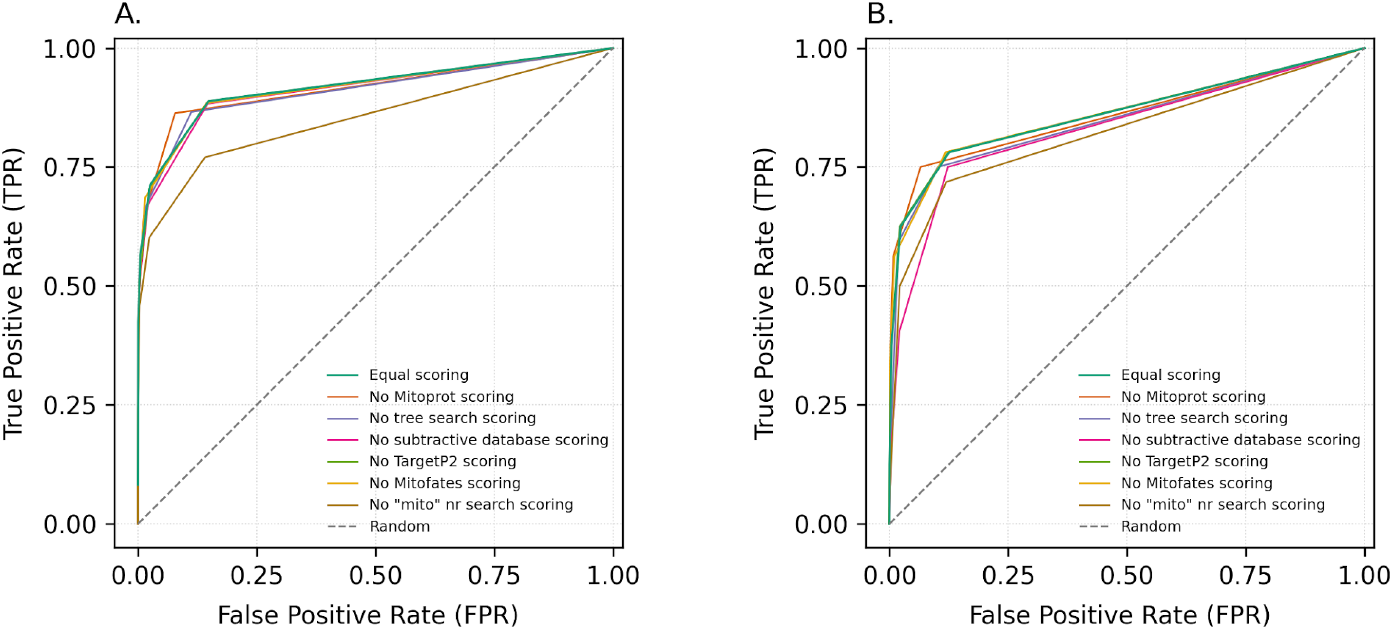
Scoring-scheme sensitivity for the *S. cerevisiae* proteome and *P. pyriformis* proteome shows that CoMR performance varies depending on component deletions. A. *S. cerevisiae* proteome. B. *P. pyriformis* proteome. These plots display ROC curves under the full set of scoring schemes, representing the effectiveness of the CoMR pipeline when using the equal scoring scheme (turquoise), when the automatic phylogenetic analysis is removed from the scoring scheme (purple), when the subtractive mitochondrial database search is removed from the scoring scheme (pink), when TargetP2 is removed from the scoring scheme (green), when Mitofates is removed from the scoring scheme (yellow), and when the NCBI non-redundant database search is removed from the scoring scheme (brown). The dashed line indicates a random classifier.

### CoMR performance on a non-model organism, *Paratrimastix pyriformis*

Because the relative contribution of individual evidence layers is likely to depend on phylogenetic context and database representation, we next evaluated CoMR performance on a divergent, non-model organism. We applied the same benchmarking framework to the anaerobic protist *Paratrimastix pyriformis*, whose spatial proteomics-derived mitochondrial-related organelle (MRO) proteome include 32 annotated proteins^35^.

Using the default equal-weight scoring scheme, we constructed a ROC curve and calculated a ROC-AUC of 0.86 for the *P. pyriformis* proteome (Figure 2B; Table 1), indicating strong discriminatory performance despite the reduced size and divergence of its MRO proteome. As observed for *S. cerevisiae*, MitoProt generated an elevated number of false positives in the *P. pyriformis dataset* (Figure 3B). Down-weighting MitoProt resulted in a modest increase in ROC-AUC (Table 1; Figure 4B), indicating that its contribution inflates false positive rates in this proteome. In contrast to S. *cerevisiae*, where NR-based homology had the largest impact on performance, both NR and SMD homology searches substantially contributed to ROC-AUC in *P. pyriformis* (Table 1; Figure 5B).

### Performance comparison between CoMR and TargetP2

Among the three targeting predictors included in CoMR, TargetP2 showed the lowest false positive rate and overall strongest standalone performance. We therefore used TargetP2 as a baseline comparator to evaluate CoMR performance on both the *S. cerevisiae* and *P. pyriformis* proteomes. For both datasets, CoMR achieved higher ROC-AUC values than TargetP2 under the equal-weight scoring scheme (Table 2; Figure 6A-B). This indicates that integration of multiple evidence layers provides improved discriminatory performance relative to targeting prediction alone.

**Table 2.** Comparing the performance of CoMR and TargetP2 on the *S. cerevisiae* and *P. pyriformis* proteomes using AUC-ROC scores.

| ROC curves | *S. cerevisiae* AUC score | *P. pyriformis* AUC score |
| --- | --- | --- |
| CoMR Equal score | 0.91782 | 0.86113 |
| TargetP2 | 0.72255 | 0.54350 |

**Figure 6.**
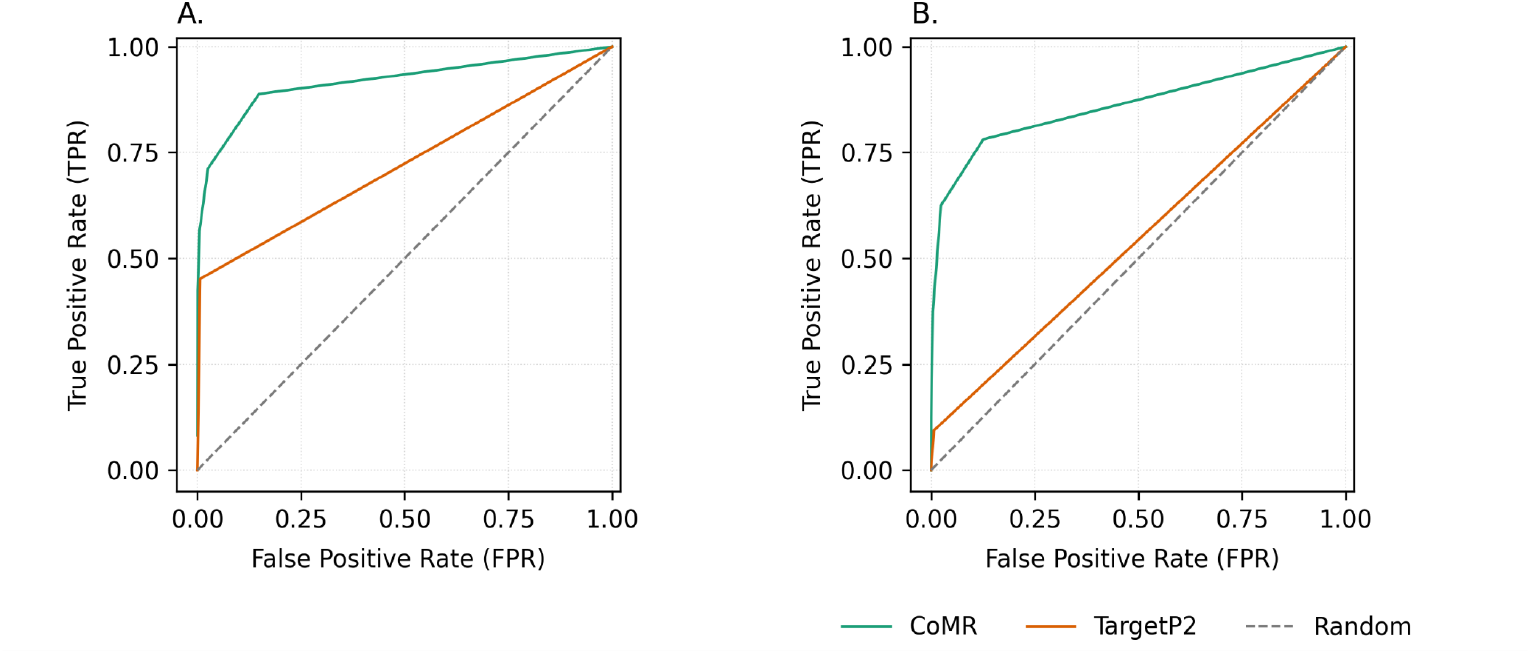
A comparison of CoMR function vs. TargetP2 using the *S. cerevisiae* and *P. pyriformis* proteomes. A. *S. cerevisiae* proteome. B. *P. pyriformis* proteome. ROC curves for CoMR using the equal scoring scheme (green), TargetP2 (orange), and a random classifier (dashed line) are displayed. A. *S. cerevisiae* proteome, B. *P. pyriformis* proteome.

Because the *P. pyriformis* dataset contains a small number of positive cases relative to the total proteome size, ROC-AUC values may overestimate predictive performance under extreme class imbalance^36^. To more directly evaluate classification precision, we calculated precision and recall across score thresholds and constructed precision-recall (PR) curves for both CoMR (equal-weight scoring scheme) and TargetP2 (Table 3, Figure 7). CoMR achieved a PR-AUC of 0.18340, substantially exceeding both the random baseline (0.00236, corresponding to the prevalence of MRO proteins in the dataset) and TargetP2 (PR-AUC = 0.01859). This represents an approximately 78-fold enrichment over random expectation and nearly a tenfold improvement over standalone targeting prediction. These results indicate that CoMR maintains strong precision-recall performance despite the extreme rarity of MRO proteins in the *P. pyriformis* proteome.

**Table 3.**
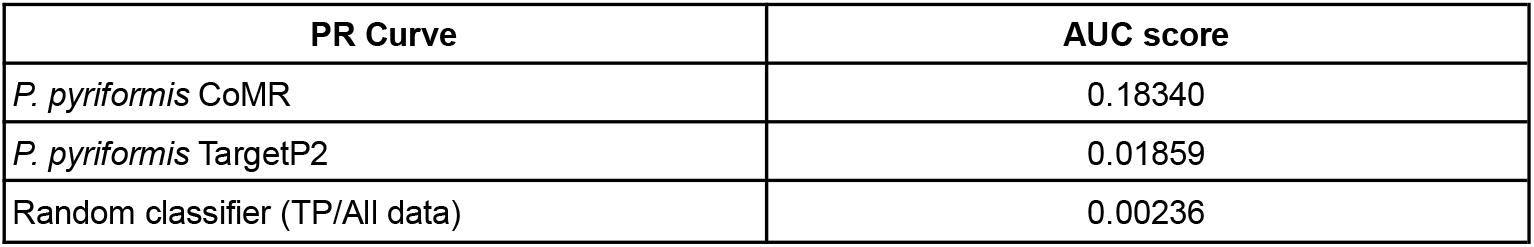
AUPR scores comparing the performance of CoMR and TargetP2 on the *P. pyriformis* proteome.

**Figure 7.**
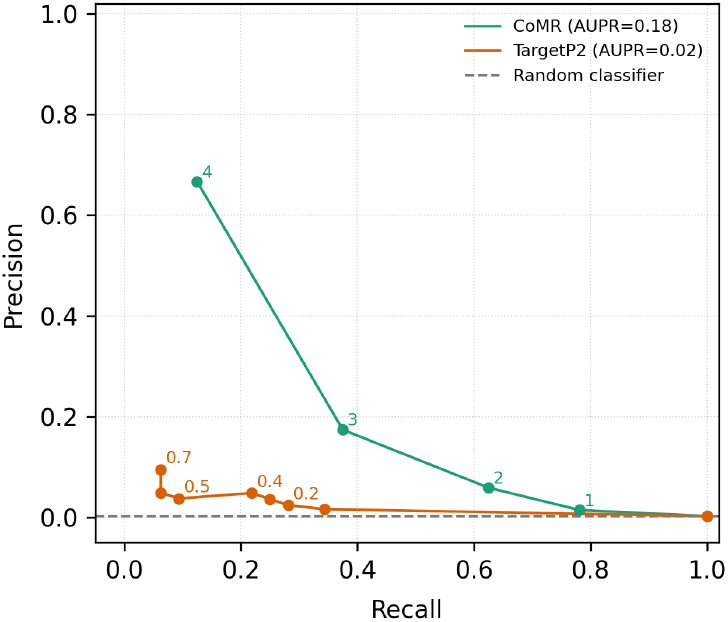
Precision-recall performance for CoMR and TargetP2 in the *P. pyriformis* proteome. PR curve for CoMR (equal scoring) and the TargetP2 PR curve (score threshold). For visualization, only thresholds yielding informative precision (i.e., non-zero, at least one true positive) are plotted. The dashed line indicates a random classifier at prevalence.

### CoMR runtime

To assess the computational cost of large-scale homology searches, we evaluated CoMR runtime and memory usage under different resource allocations and with or without NR searches enabled (Table 4). Under high-performance settings (128 CPUs; DIAMOND threads = 64; block size = 4), complete execution required 3 h 12 min for *S. cerevisiae* (5,967 proteins) and 5 h 11 min for *P. pyriformis* (13,532 proteins), with peak memory usage of 200 GB and 228 GB, respectively. Reducing computational allocation (24 CPUs; DIAMOND threads = 8; block size = 1) increased runtime and reduced memory consumption. For *S. cerevisiae*, runtime increased from 3 h 12 min to 7 h 26 min, while peak RAM decreased from 200 GB to 88 GB. A similar scaling trend was observed for *P. pyriformis*, with runtime increasing from 5 h 11 min to 10 h 27 min and peak RAM decreasing from 228 GB to 115 GB.

**Table 4.**
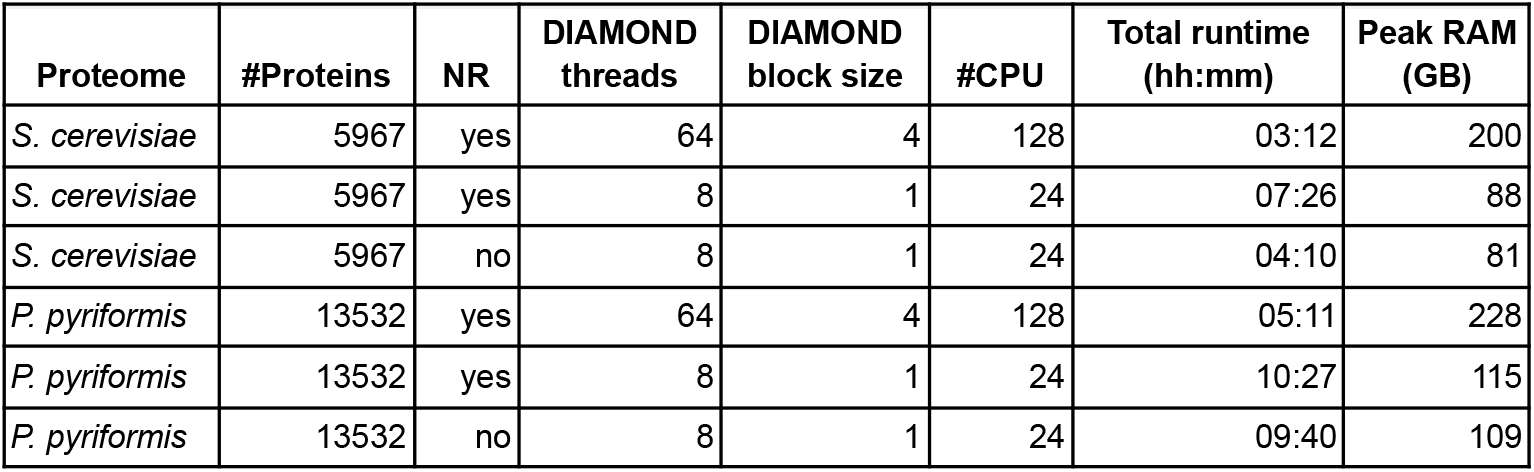
Computational performance of CoMR across hardware configurations. CoMR was executed on the *Saccharomyces cerevisiae* (5,967 proteins) and *Paratrimastix pyriformis* (13,532 proteins) proteomes under two hardware configurations: a high-performance computing cluster and under a reduced-performance allocation. The table reports the number of DIAMOND threads and block size used for the NR homology search, the total number of CPU cores allocated, total runtime, and peak memory usage (GB). Runtime corresponds to the complete pipeline execution including targeting prediction, homology searches, phylogenetic reconstruction, and score integration.

Disabling NR searches under reduced-resource settings moderately reduced runtime and memory usage. In *S. cerevisiae*, runtime decreased from 7 h 26 min to 4 h 10 min when NR was disabled, while peak RAM decreased from 88 GB to 81 GB. In *P. pyriformis*, runtime decreased from 10 h 27 min to 9 h 40 min and peak RAM from 115 GB to 109 GB. Although NR searches represent a major computational component, substantial runtime persisted even when NR was disabled, particularly for the larger *P. pyriformis* proteome. This likely reflects the computational cost of multiple sequence alignment and phylogenetic tree reconstruction steps, which scale with the number of candidate protein families identified.

## Discussion

Mitochondrial proteome reconstruction remains challenging, particularly in phylogenetically diverse or poorly represented eukaryotic lineages. Here, we introduce CoMR, an integrative workflow that combines targeting prediction, curated homology searches, large-scale NR searches, and automated phylogenetic analysis into a unified scoring framework. Across both a well-annotated model organism (*Saccharomyces cerevisiae*) and a divergent anaerobic protist (*Paratrimastix pyriformis*), CoMR demonstrated strong discriminatory performance and consistent improvement over standalone targeting prediction.

### Integrative scoring improves over targeting prediction alone

In *S. cerevisiae*, CoMR achieved an ROC-AUC of 0.92, substantially outperforming TargetP2 (ROC-AUC = 0.72). This difference underscores a central finding of this study: mitochondrial targeting signals alone are insufficient for comprehensive proteome reconstruction. While targeting predictors performed well in minimizing false negatives, their false positive rates varied considerably, particularly for MitoProt. The modest changes observed when altering MitoProt weighting demonstrate that CoMR is robust to individual predictor biases, likely because independent evidence layers buffer one another. This integrative behavior is a key advantage over single-method approaches.

### Evidence-layer contributions are lineage dependent

Ablation analyses revealed that the relative importance of evidence layers varies across phylogenetic contexts. In *S. cerevisiae*, removal of NR-based homology had the largest impact on ROC-AUC, illustrating the dense representation of fungal sequences in public databases. In contrast, in *P. pyriformis*, both NR and curated SMD homology contributed substantially to predictive performance. This shift likely reflects reduced representation of metamonad sequences in NR and highlights the value of curated mitochondrial reference datasets in poorly sampled lineages. These results emphasize that no single evidence type is universally dominant. Instead, optimal mitochondrial prediction depends on phylogenetic context and database coverage. The ability of CoMR to integrate multiple partially independent signals appears particularly beneficial in non-model organisms.

### Performance under extreme class imbalance

The *P. pyriformis* dataset presents a highly imbalanced classification problem, with only 32 annotated MRO proteins among more than 13,000 sequences. While ROC-AUC values remained strong (0.86), precision-recall analysis provided a more informative assessment. CoMR achieved a PR-AUC of 0.183, representing a ∼78-fold enrichment over random expectation and nearly a tenfold improvement over TargetP2. These results demonstrate that CoMR maintains meaningful precision even when true positives are rare, an essential property for mitochondrial reconstruction in reduced or highly derived organelles.

### Robustness and customizability of the scoring framework

Down-weighting or removing individual evidence layers resulted in only modest changes in performance, suggesting that CoMR is not overly dependent on any single signal. This robustness supports the use of the default equal-weight scoring scheme across diverse datasets. At the same time, the customizable scorecard architecture allows users to adjust component weights to reflect lineage-specific knowledge or data availability. Such flexibility may be particularly valuable when investigating deeply branching or sparsely sampled eukaryotic groups.

### Limitations

Several limitations should be acknowledged. First, benchmarking relied on curated proteomes, which may themselves be incomplete or subject to annotation bias. Second, NR-based evidence is inherently influenced by database composition; although taxonomic exclusion was implemented to prevent circularity, residual biases toward well-represented lineages remain unavoidable. Third, automated phylogenetic placement relies on reference alignment quality and may be less reliable for highly divergent proteins.

Future improvements could incorporate additional lineage-specific reference datasets. Experimental validation in additional non-model systems would further strengthen generalizability.

## Conclusions

Together, these results demonstrate that integrative evidence scoring substantially improves mitochondrial proteome reconstruction over standalone targeting prediction, particularly in non-model and database-sparse lineages. CoMR provides a reproducible and flexible framework for mitochondrial and mitochondrial-related organelle prediction, enabling systematic comparative analyses across diverse eukaryotes.

## Resources availability

### Data and code availability

The CoMR workflow, including all scripts, configuration templates, and Snakemake rules, is publicly available via the CoMR GitHub repository (https://github.com/theLabUpstairs/CoMR). CoMR databases and benchmarking-modified databases and associated scripts are publicly available via the CoMR Figshare repository (https://doi.org/10.17044/scilifelab.31361839). All software dependencies are containerized, and reproducible execution is supported via Docker (ghcr.io/thelabupstairs/comr:git-2bc33bf) or Singularity/Apptainer image, available from the CoMR Figshare repository (https://doi.org/10.17044/scilifelab.31361839). Documentation, installation instructions, and usage examples are provided within the CoMR github repository, including system-specific execution guides and workflow descriptions. These resources are intended to support reproducible deployment and adaptation of the pipeline to diverse computational environments.

## Key points

- CoMR is an integrative scoring pipeline that combines targeting prediction, curated and large-scale homology searches, and automated phylogenetic analysis for mitochondrial proteome reconstruction.
- In *Saccharomyces cerevisiae*, CoMR achieved strong discriminatory performance (ROC-AUC = 0.92), outperforming standalone TargetP2 prediction (ROC-AUC = 0.72).
- In the divergent metamonad *Paratrimastix pyriformis*, CoMR maintained robust performance (ROC-AUC = 0.86; PR-AUC = 0.183) despite extreme class imbalance.
- Ablation analyses demonstrate that the relative contribution of evidence layers is lineage-dependent, highlighting the importance of integrative approaches in poorly represented clades.
- CoMR provides a reproducible and scalable framework applicable to both model and non-model eukaryotes.

## Acknowledgments

The authors would like to thank Viktor Törnblom for technical assistance and for testing the CoMR pipeline during development.

