## Supplementary Material for "CoMR: an integrative scoring pipeline for Comprehensive Mitochondrial proteome Reconstruction across eukaryotes"

### Supplementary Data

#### Supplementary Methods - CoMR databases

##### CoMR's Subtractive Mitochondrial Database.

The SMD is derived from well-curated whole-cell proteomes and mitochondrial proteomes of six model organisms: *Andalucia godoyi*, *Tetrahymena thermophila*, *Arabidopsis thaliana*, *Homo sapiens*, *Acanthamoeba castellanii*, *Saccharomyces cerevisiae*. For *Andalucia godoyi*, mitochondrial proteins were defined via a comprehensive *in silico* approach applied to the draft nuclear genome (GenBank accession VRVR00000000), which identified 864 candidate mitochondrial proteins based on homology and functional annotation consistent with established mitochondrial protein families[1]. Although derived from genome-based annotation rather than organelle proteomics, *Andalucia godoyi* was included to increase the phylogenetic diversity of the reference set by representing a deeply branching jakobid lineage. For *Tetrahymena thermophila*, the mitochondrial protein set was derived from highly purified mitochondrial fractions subjected to exhaustive tryptic digestion followed by tandem liquid chromatography-tandem mass spectrometry (LC-LC-MS/MS), resulting in direct identification of mitochondrial proteins[2,3] (predicted protein Genbank accession EAR80512 to EAS07932). For *Arabidopsis thaliana*, mitochondrial assignments were based on proteomic analyses of purified mitochondria using two-dimensional electrophoresis and LC-MS/MS, integrated with genome-based annotation to confirm protein identities and subcellular localization[4] (NCBI Bioproject PRJNA10719 and SAMN03081427; The Arabidopsis Genome Initiative, 2000). The human mitochondrial proteome was retrieved from UniProt entries annotated with GO term GO:0005739 (mitochondrion), restricted to reviewed entries with experimental or curated evidence supporting subcellular localization (International Human Genome Sequencing Consortium, 2001[5]; mitochondrial proteins from UniProt (GO:0005739)). For *Acanthamoeba castellanii*[6,7], the mitochondrial proteome was established through a combined proteomic and bioinformatic approach, including mass spectrometry analysis of mitochondrial fractions and subsequent curation against genomic and transcriptomic data to validate protein assignments (NCBI Bioproject PRJNA599339 and PRJNA487265). The *Saccharomyces cerevisiae* mitochondrial set was based on curated annotations from the Saccharomyces Genome Database (SGD), integrating genetic, biochemical, and proteomic evidence accumulated through community curation[8] (https://www.yeastgenome.org/). For each of the six organisms, each sequence in the mitochondrial proteome was used as a query in a Diamond search of the whole cell proteome, using the "more-sensitive" setting and an e-value cutoff of 0.0001. We collected the hits from the whole cell proteome that were >90% identical to a mitochondrial query on the amino acid level and removed these hits from the whole cell proteome. This generated two datasets for each of the six organisms: a dataset containing mitochondrial and mitochondrial-like proteins, and a dataset containing proteins <90% identical to a mitochondrial protein in each organism. The SMD consists of all twelve datasets generated above in a two FASTA formatted file (SubstractiveDB and MitoDB) with headers that describe their mitochondrial or non-mitochondrial designation. For the CoMR pipeline, each sequence from the input set of protein sequences is used as a query in a Diamond search against the SMD, the results of which are then filtered by their bit score. All CoMR databases are available on the CoMR Figshare repository (https://doi.org/10.17044/scilifelab.31361839).

#### Phylogenetic reconstruction and parsing.

We constructed HMM profiles based on the SMD by clustering the proteomes of all six organisms using Broccoli v1.2[9] to create orthologous groups (OGs), which were then manually curated. These OGs contain proteins with headers signifying if they belong to a mitochondrial or non-mitochondrial dataset within the SMD. These OGs were then aligned with mafft v7.310 einsi[10], trimmed with trimal v1.4.rev15[11] using the -gappyout setting, and the resulting alignments were used to construct HMM profiles using HMMER 3.3.2[12] with default settings. For the CoMR pipeline, each sequence in the input set of protein sequences is searched against a database of these HMM profiles with HMMscan. If a hit is identified that input sequence is then added to the corresponding untrimmed alignment that produced the HMM profile. If more than one hit is identified, the input sequence is added to alignment with the HMM profile that has the highest scoring hit based on its e-value. These sequences are then aligned and trimmed using the methods mentioned above, and used to estimate a phylogeny with IQTree 2[13] with the LG+gamma model of evolution. Each phylogeny is parsed by rooting the tree at the input sequence in question, and the input sequence is assessed for its relationship to other sequences in the tree that were either mitochondrial or non-mitochondrial. The Fitch parsimony algorithm for ancestral reconstruction of states was then used to predict whether the query sequence was “mitochondrial” (MITO), or “non-mitochondrial” (OTHER)[14].

### Supplementary Methods - CoMR evidence integration and scoring

The CoMR composite score integrates six independent evidence layers reflecting support for mitochondrial localization: (i) TargetP2[15], (ii) MitoProt II[16], (iii) MitoFates[17], (iv) curated homology support from the Subtractive Mitochondrial Database (SMD), (v) large-scale homology support from the NCBI non-redundant (NR) database, and (vi) phylogenetic placement based on automated tree parsing.

Each evidence layer is converted into a binary signal prior to score integration. TargetP predictions are considered positive when the predicted localization class is mTP. MitoProt predictions are considered positive when the export probability is ≥70%. MitoFates predictions are considered positive when the sequence is classified as “Possessing mitochondrial presequence”. Curated homology support is considered positive when a sequence has top hits to three or more mitochondrial datasets within the SMD. NR-based homology support is considered positive when at least one retained DIAMOND[18] hit is annotated with a mitochondrion-related keyword after taxonomic exclusion. Phylogenetic support is considered positive when automated tree parsing assigns the query sequence to a mitochondrial clade.

DeepMito[15] predictions are not included in the composite score, as this method assumes mitochondrial localization of all input sequences and instead predicts sub-mitochondrial localization. DeepMito outputs are reported alongside the composite score for contextual interpretation. Under the default “equal” scoring scheme, each positive evidence layer contributes one point to the composite score. The resulting CoMR score ranges from 0 (no supporting evidence) to 6 (support from all evidence layers).

### Supplementary Methods - CoMR performance evaluation

#### Reconstruction of reference databases for benchmarking

To prevent circularity during benchmarking on *Saccharomyces cerevisiae*, we generated benchmarking-specific versions of the internal CoMR reference databases in which *S. cerevisiae* sequences were excluded. These filtered databases were used exclusively for benchmarking analyses and were not employed for standard CoMR runs. All CoMR benchmarking-specific databases are available on the CoMR Figshare repository (https://doi.org/10.17044/scilifelab.31361839).

#### Filtered Subtractive Mitochondrial Database (SMD)

For benchmarking on *S. cerevisiae,* a filtered version of the Subtractive Mitochondrial Database (SMD) was constructed by removing all *S. cerevisiae* sequences from the precomputed SMD FASTA files. In the original SMD, *S. cerevisiae* sequences were identified by a `YEAST` prefix in the FASTA headers, as well as by explicit *Saccharomyces cerevisiae* annotations. All sequences whose headers contained either the `YEAST` prefix or Saccharomyces cerevisiae identifiers were removed. The resulting filtered SMD was used to rebuild DIAMOND databases for homology searches during benchmarking. No other modifications were applied to the SMD.

#### Filtered orthologous group alignments and HMM profiles

Benchmarking-specific orthologous group (OG) alignments were generated by filtering the original SMD-derived multiple sequence alignments to remove all sequences labeled as *S. cerevisiae* using the same header-based identifiers. After filtering, OGs with zero sequences were discarded. No OGs were reduced to a single sequence; all retained OGs contained at least two sequences and were realigned with mafft v7.526, trimmed with trimal v1.5.1 using the -gappyout setting, and used to reconstruct hidden Markov model (HMM) profiles using HMMER v3.3.2 with default parameters. The resulting HMM profiles were concatenated and indexed to produce a benchmarking-specific HMM database, which was used for all HMM-based searches during *S. cerevisiae* benchmarking.

#### NR homology search parsing for benchmarking

To assess the contribution of large-scale homology searches during benchmarking and to prevent circularity, DIAMOND searches against the NCBI non-redundant (NR) protein database were parsed using a taxonomically controlled filtering strategy. DIAMOND searches were performed with taxonomy reporting enabled, such that taxonomic identifiers (staxids), scientific names, and kingdom-level annotations were included in the output for each hit.

Parsed NR results were processed on a per-query basis, and taxonomic filtering was applied prior to hit ranking and evidence assignment. For each query protein, all DIAMOND hits assigned to excluded taxonomic identifiers were removed before identifying top hits, non-hypothetical hits, or mitochondrial-related hits. If no hits remained after filtering, the query was retained in the parsed output with all NR-related fields recorded as “None,” ensuring consistent downstream integration with other evidence layers.

#### Taxonomic exclusion strategies

Because entries in NR may be annotated with taxonomic identifiers corresponding to strains, isolates, or assemblies rather than the canonical species-level taxid, taxonomic exclusion was implemented by removing all descendant taxonomic identifiers of a specified root node in the NCBI taxonomy[19]. Descendant taxids were identified using the NCBI taxonomy hierarchy (nodes.dmp), ensuring that all strain- and isolate-level annotations associated with a given taxon were excluded.

Three exclusion strategies were applied during benchmarking of the *Saccharomyces cerevisiae* proteome:

##### *Species-level exclusion*

All DIAMOND hits assigned to *S. cerevisiae* (taxid 4932) and its taxonomic descendants were excluded. This strategy prevents direct self-hits while retaining closely related fungal homologs, reflecting a realistic benchmarking scenario for a well-studied model organism.

##### *Genus-level exclusion*

All hits assigned to any species within the genus *Saccharomyces* and their descendants were excluded. This stricter condition reduces the contribution of very closely related homologs and evaluates the robustness of NR-based evidence when immediate phylogenetic neighbors are removed.

##### *Order-level exclusion*

All hits assigned to members of the corresponding fungal order and their descendants were excluded. This exclusion represents a conservative scenario approximating benchmarking conditions for non-model organisms with limited representation in NR.

For each exclusion strategy, the same DIAMOND output was parsed independently using the corresponding exclusion set, without rerunning the homology searches.

For *Paratrimastix pyriformis*, all DIAMOND hits assigned its NCBI Taxonomy ID (taxid 342808, no descendant) were excluded.

#### Reproducibility and robustness

All taxonomic exclusions were implemented deterministically using explicit taxonomic identifiers derived from the NCBI taxonomy hierarchy, ensuring reproducibility across runs. The three exclusion levels for *S. cerevisiae* were used to assess the sensitivity of CoMR performance to the availability of closely related homologs in NR, while preserving identical inputs, scoring schemes, and downstream analyses across benchmarking runs.

#### Statistical analysis for benchmarking

CoMR benchmarking was performed on both *S. cerevisiae* and *P. pyriformis* by transforming their own CoMR scored CSV outputs into binary evaluation tables, computing threshold-based performance metrics, and deriving ROC-AUC values.

First, for each CoMR scored CSV, an evaluation table was constructed containing for each sequence, its CoMR Score as well as the actual classification from literature (mitochondrial or not mitochondrial). Threshold-based benchmarking was then carried out for score cutoffs S = 0-6 (inclusive). At each threshold, sequences with CoMR_Score ≥ S were considered positive predictions and used to compute TP, FP, TN, and FN counts, from which TPR, FPR, and precision were calculated.

ROC-AUC values were computed by sorting ROC points by FPR and integrating the TPR-FPR curve. The default integration used the trapezoidal rule (linear interpolation between adjacent ROC points). For comparison, a left-rectangle (step) approximation is also supported, and the per-segment decomposition into rectangle and triangular correction terms is reported alongside the corrected segment areas and overall AUC.

The Equal scoring scheme uses the standard CoMR scorecard where each component (subtractiveDB, targetp, mitoprot, mitofates, mito nr search, tree) contributes equally (value = 1; customDB = 0), and default thresholds are applied. ROC points are computed for score thresholds 0-6, reporting true-positive rate (TPR) and false-positive rate (FPR) at each threshold. Curves are plotted along with a random-classifier baseline (diagonal). For scoring-scheme comparisons, CoMR scores were recomputed from the scored CSVs using the same component rules with scheme-specific changes to component weights (removing TargetP2, removing Mitoprot, downweighting Mitoprot by half, removing Mitofates, removing subtractiveDB, removing Trees, removing NR search). ROC points and AUC were then derived for each scheme and dataset. ROC curves were generated from threshold-based metrics computed on the CoMR scored CSV outputs. A random-classifier baseline (diagonal) was plotted for reference.

We evaluated the three MTS predictors (TargetP, MitoProt, and MitoFates) by computing confusion counts at each predictor's recommended cutoff implemented in CoMR. TargetP predictions were considered positive when tp_prediction = mTP, MitoProt predictions when mp_score ≥ 70, and MitoFates predictions when mf_seq = Possessing_mitochondrial_presequence. For each predictor, we counted true positives (TP), false positives (FP), true negatives (TN), and false negatives (FN). For TargetP/CoMR comparison, CoMR ROC curves were taken from the equal-scoring scheme derived from thresholds 0-6. TargetP2 ROC points were computed using the recommended TargetP2 cutoff (tp_prediction = mTP) on the evaluation tables. To enable ROC-AUC calculation for the binary TargetP2 classifier, endpoints (0,0) and (1,1) were included. AUC values were computed by trapezoidal integration of the TPR-FPR curves. A random-classifier baseline (diagonal) was plotted for reference.

Precision-recall curves were derived for the *P. pyriformis* benchmark using the equal-scoring CoMR thresholds (0-6) and TargetP2 probability thresholds (0.0-1.0 in 0.1 increments) based on tp_score from the scored CSV. Thresholds that did not change the operating point (CoMR 5-6, TargetP2 0.8-1.0) were excluded from plotting for clarity. Precision and recall were computed from TP, FP, and FN counts; CoMR values were taken from the ROC points table, and TargetP2 values were computed from the scored CSV. All thresholds are retained in both the plot and the output table, which also reports AUPR per predictor. A random-classifier baseline was plotted at the prevalence (TP / all sequences), and the random AUPR equals this prevalence.

All CoMR benchmarking-specific data and associated scripts are available on the CoMR Figshare repository (https://doi.org/10.17044/scilifelab.31361839).

#### Supplementary Figures and Tables

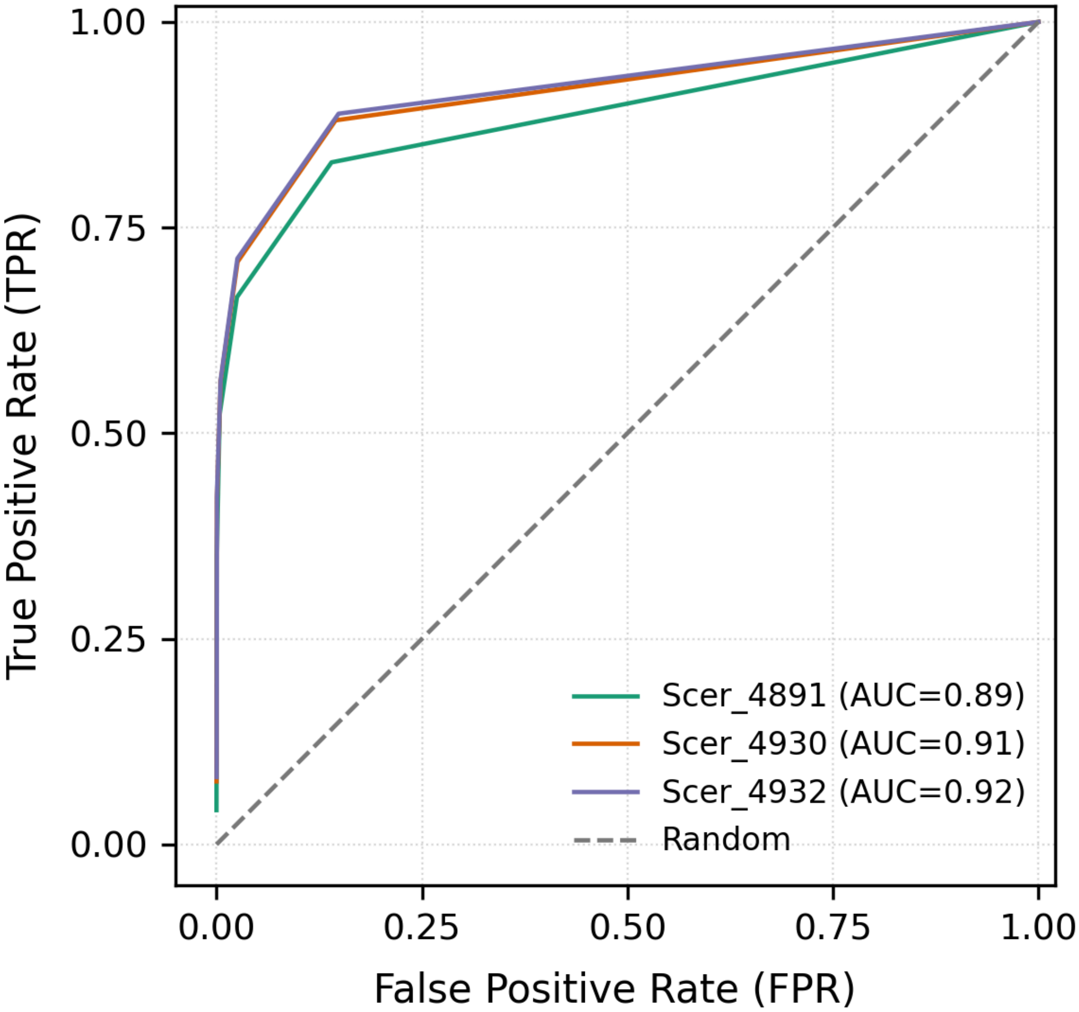


**Figure S1: The ROC curves for the *S. cerevisiae proteome* classified by CoMR indicate robust discrimination across benchmarking runs.**
ROC curves for all yeast benchmark runs under the Equal scoring scheme. Each colored line represents one run and is annotated with its AUC; the dashed line indicates a random classifier. The runs correspond to taxonomic exclusion levels for *S. cerevisiae* benchmarking (species *S. cerevisiae* (taxid 4932 and descendants); genus *Saccharomyces* (taxid 4930 and descendants); order Saccharomycetales (taxid 4891 and descendants). Axes show false-positive rate (FPR) and true-positive rate (TPR).

**Table S1. AUC scores of ROC curves summarizing CoMR performance on *S. cerevisiae* according to different scoring schemes and taxonomic exclusion levels.**

| Scoring scheme | Taxonomic exclusion level | *S. cerevisiae* AUC score |
|---|---:|---:|
| Equal scoring | Species (taxid: 4932) | 0.91782 |
| No Mitoprot scoring | Species (taxid: 4932) | 0.91482 |
| Mitoprot downweighted | Species (taxid: 4932) | 0.92202 |
| No tree search scoring | Species (taxid: 4932) | 0.91074 |
| No subtractive database scoring | Species (taxid: 4932) | 0.91285 |
| No TargetP2 scoring | Species (taxid: 4932) | 0.91736 |
| No Mitofates scoring | Species (taxid: 4932) | 0.91613 |
| No "mito" nr search scoring | Species (taxid: 4932) | 0.85265 |
| Equal scoring | Genus (taxid: 4930) | 0.91410 |
| No Mitoprot scoring | Genus (taxid: 4930) | 0.90936 |
| Mitoprot downweighted | Genus (taxid: 4930) | 0.91793 |
| No tree search scoring | Genus (taxid: 4930) | 0.90731 |
| No subtractive database scoring | Genus (taxid: 4930) | 0.90871 |
| No TargetP2 scoring | Genus (taxid: 4930) | 0.91360 |
| No Mitofates scoring | Genus (taxid: 4930) | 0.91231 |
| No "mito" nr search scoring | Genus (taxid: 4930) | 0.85316 |
| Equal scoring | Order (taxid: 4891) | 0.88672 |
| No Mitoprot scoring | Order (taxid: 4891) | 0.87452 |
| Mitoprot downweighted | Order (taxid: 4891) | 0.88973 |
| No tree search scoring | Order (taxid: 4891) | 0.87297 |
| No subtractive database scoring | Order (taxid: 4891) | 0.87810 |
| No TargetP2 scoring | Order (taxid: 4891) | 0.88555 |
| No Mitofates scoring | Order (taxid: 4891) | 0.88204 |
| No "mito" nr search scoring | Order (taxid: 4891) | 0.85315 |
